# Linking White-Matter Development to Clinical Variation in Autism: Longitudinal Normative Modelling of Fractional Anisotropy (FA) in the EU-AIMS LEAP Cohort

**DOI:** 10.1101/2025.11.25.690402

**Authors:** Ramona Cirstian, Natalie J. Forde, Barbora Rehák Bučková, Flavio Dell’Acqua, Tony Charman, Eva Loth, Beth Oakley, Laura Colomar Mollà, Isabel Yorke, Richard Stones, Declan G. M. Murphy, Tobias Banaschewski, Jan K. Buitelaar, EU-AIMS LEAP Group, Andre F. Marquand, Christian F. Beckmann

**Author notes:** Contributed equally as senior authors (Andre F. Marquand, Christian F. Beckmann).

## Abstract

**Background:** Autism is characterised by neurobiological and clinical heterogeneity, which often limits the sensitivity of traditional group comparisons. Longitudinal normative modelling offers a way to capture heterogeneity and to describe an individual’s brain–behaviour relationships beyond diagnostic categories.

**Methods:** We analysed diffusion MRI scans from 3 waves of the EU-AIMS Longitudinal European Autism Project (LEAP). Using normative models of white matter fractional anisotropy (FA) trained on a neurotypical reference cohort, we computed individual deviations relative to age-typical norms. Our sample included 544 individuals (Wave 1: 189 autistic 162 non-autistic; Wave 2: 162 autistic136 non-autistic; Wave 3: 145 autistic 84 non-autistic), aged 7–37 years, with mean intervals of ∼1.5 years (W1–W2) and ∼6 years (W2–W3). We first tested for group level differences between autistic and non-autistic participants in FA deviations across 48 white matter tracts and then examined the associations between these deviation scores and sensory, adaptive, cognitive, and behavioural measures across development within the autism group.

**Results:** Group-wise comparisons revealed no significant differences in FA deviations between autistic and non-autistic participants. Within the autism group, multivariate analysis (CCA) showed significant (p<0.05) cross-sectional brain-behaviour associations at Wave 1 (r=0.26) and Wave 2 (r=0.32), but not at Wave 3. Longitudinally, we found significant multivariate associations only for the W1–W3 interval (r=0.36). Follow-up univariate correlations supported these findings, with Wave 2 showing widespread moderate (r≈0.35) associations while longitudinally, the univariate associations were generally weaker (r≈0.2). Across analyses, higher FA was generally associated with higher Vineland Adaptive Behaviour Scales (VABS) and lower Short Sensory Profile (SSP) scores. Associations with the Autism Diagnostic Observation Schedule (ADOS) were mixed, and the Social Responsiveness Scale (SRS) and Repetitive Behaviour Scale (RBS) showed wave-dependent relationships, with specific tract involvement varying across timepoints.

**Conclusions:** White-matter organisation in autism does not differ from that of non-autistic on average but rather follows a dimensional architecture within the autism phenotype. We showed that person-specific deviations from shared neurodevelopmental pathways are linked to behavioural variation among autistic individuals, which cannot be captured by group-level approaches alone. Longitudinal normative modelling provides a practical approach to understand and quantify this heterogeneity.

**EU-AIMS LEAP Group:** Zuzana Suchomelova, Mee Rim Oh, Chirag Mehra, Mei Lin Law, Sanjana Gandhi, Laura Bravo Balsa, Maria Dauvermann, Isabel Yorke, Beth Oakley, Rosemary J. Holt, Edward Bullock, Yumnah Kahn, Esme Hayes, Leona Strauss, Noel Lam, Anouk Dykstra, Kim Lamers, Marije Mars, Lucas Geelen, Lotte Beckers, Anna Praat, Feline van Aagten, Sjors Reith, Viola Hollestein, Sanne Kluin, Natalie Forde, Jill Naaijen, Anna Kaiser, Sarah Baumeister, Pascal-M. Aggensteiner, Jumana Ahmad, Sara Ambrosino, Bonnie Auyeung, Tobias Banaschewski, Simon Baron-Cohen, Sarah Baumeister, Christian F. Beckmann, Sven Bölte, Thomas Bourgeron, Carsten Bours, Michael Brammer, Daniel Brandeis, Claudia Brogna, Yvette de Bruijn, Jan K. Buitelaar, Bhismadev Chakrabarti, Tony Charman, Ineke Cornelissen, Daisy Crawley, Flavio Dell’Acqua, Guillaume Dumas, Sarah Durston, Christine Ecker, Jessica Faulkner, Vincent Frouin, Pilar Garcés, David Goyard, Lindsay Ham, Hannah Hayward, Joerg Hipp, Rosemary J. Holt, Mark H. Johnson, Emily J. H. Jones, Prantik Kundu, Meng-Chuan Lai, Xavier Liogier D’ardhuy, Michael V. Lombardo, Eva Loth, David J. Lythgoe, René Mandl, Andre Marquand, Luke Mason, Maarten Mennes, Andreas Meyer-Lindenberg, Carolin Moessnang, Nico Mueller, Declan G. M. Murphy, Laurence O’Dwyer, Marianne Oldehinkel, Bob Oranje, Gahan Pandina, Antonio M. Persico, Jack Price, Annika Rausch, Barbara Ruggeri, Amber N. V. Ruigrok, Jessica Sabet, Roberto Sacco, Antonia San Jóse Cáceres, Emily Simonoff, Will Spooren, Julian Tillmann, Roberto Toro, Heike Tost, Jack Waldman, Steve C. R. Williams, Caroline Wooldridge, Iva Ilioska, Ting Mei, Marcel P. Zwiers.

**Key Points:** *Question:* In the context of heterogeneity in autism, what do individual differences in white-matter development reveal about brain–behaviour relationships over time?

*Findings:* In this study, we found significant relationships between individual white-matter deviations and clinical traits in autism both cross-sectionally as well as longitudinally. Increased adaptive functioning and decreased sensory processing were associated with higher FA in long-association and commissural pathways, including the cingulum bundle, superior longitudinal fasciculus, and corpus-callosum, with longitudinal effects most evident over a 7.5-year interval.

*Meaning:* Developmentally informed normative modelling of white matter offers a pathway toward biologically grounded stratification for precision medicine in autism.

## 1. Introduction

Autism represents a broad and complex pattern of neurodevelopmental variation affecting communication, behaviour and cognition. First described by Kanner in 1943 as what he termed a “disturbance in human connectedness”, its interpretation evolved through theories characterising autism from a spectrum of social and communicative differences, distinct cognitive processing styles and difficulties in understanding others’ emotions.^1–4^ These perspectives led to the current DSM-5 definition, which recognises autism as representing a spectrum of diverse social and behavioural characteristics.^5^ It is increasingly evident from studies in both neuroimaging and genetics that behavioural definitions only partially describe autism, which reflects a combination of interacting mechanisms rather than a single defining feature.^6^

The study of white matter has helped illustrate how abnormal brain connections may be related to various behavioural and cognitive traits in autistic individuals.^7^ Diffusion MRI is a non-invasive technique used to characterise white matter fibres by measuring the movement of water molecules within tissue.^8,9^ In this context, diffusion tensor imaging (DTI) quantitatively assesses the directional movement of water, specifically fractional anisotropy (FA), which serves as a measure of microstructural organisation.^10,11^ Cross-sectional diffusion MRI studies report altered FA across association, commissural, and language-related fibre tracts in autism, while also highlighting methodological and developmental variability, supporting white matter as a relevant substrate for studying autistic heterogeneity.^12–15^

Although autism is considered relatively stable across the lifespan, the way it is behaviourally expressed and functionally manifested changes across development. Autistic characteristics, adaptive functioning, and mental health can change from childhood through adulthood, with long-term studies reporting variability in diagnostic stability and clinical trajectory.^16–19^ Large-scale projects such as the Longitudinal European Autism Project (LEAP) combine clinical, cognitive, and neuroimaging data across the lifespan, showing in recent work that individual differences in structural and functional brain organisation are linked to dimensional variation in autistic traits.^20–22^ Existing longitudinal studies using structural MRI reveal atypical cortical thinning and non-linear volumetric trajectories in autistic participants relative to non-autistic participants, while functional imaging has shown altered maturation of key networks involved in social and cognitive function.^23–28^ In contrast, longitudinal diffusion imaging studies are scarce, with only a few studies linking changes in white-matter to differences in core autism-related features.^29,30^ These findings point to a critical gap in understanding white matter development in autism and its link to clinical features.

Capturing the diversity of developmental trajectories in autism remains a challenge for neuroimaging research. Normative modelling addresses this challenge by quantifying individual brain measure deviations from typical developmental patterns.^31,32^ By considering confounding factors such as age, sex, and site variation, these models estimate typical brain-behaviour relationships across multi-site neuroimaging data.^33–36^ Normative models have been used to characterise the diversity of brain structure in autism and other psychiatric and neurodegenerative conditions.^37–41^

The goal of this study is to characterise individual differences in white-matter development in autism using longitudinal diffusion MRI data from LEAP. Specifically, we aim to (i) quantify individual deviations in white-matter microstructure using diffusion-based normative models of FA within the LEAP cohort; (ii) test whether these deviations differ between the autism and no-autism groups or relate to clinical features within the autism group; and to (iii) assess longitudinal changes in white-matter across LEAP’s multiple timepoints, evaluating whether these changes in white matter relate to changes in clinical measures over time. Together, these aims provide a longitudinal framework for linking white-matter organisation with clinical heterogeneity in autism.

Based on the considerable heterogeneity of autism, we expected modest, spatially variable associations between white-matter organisation and clinically relevant features, including core social-communication difficulties, repetitive behaviours, and adaptive functioning. We hypothesised that deviation scores would provide an individual-level framework for characterising changes in brain–behaviour relationships over time. If large effects were present, we anticipated that these would be concentrated in long association and commissural tracts previously implicated in autism (e.g. superior longitudinal fasciculus, corpus callosum).^12,42^ Overall, we expected that the main differences in brain–behaviour associations would emerge at the level of individual white-matter developmental trajectories rather than in static cross-sectional group comparisons.

## 2. Methods

An overview of our workflow is provided in Figure 1.

**Figure 1.**
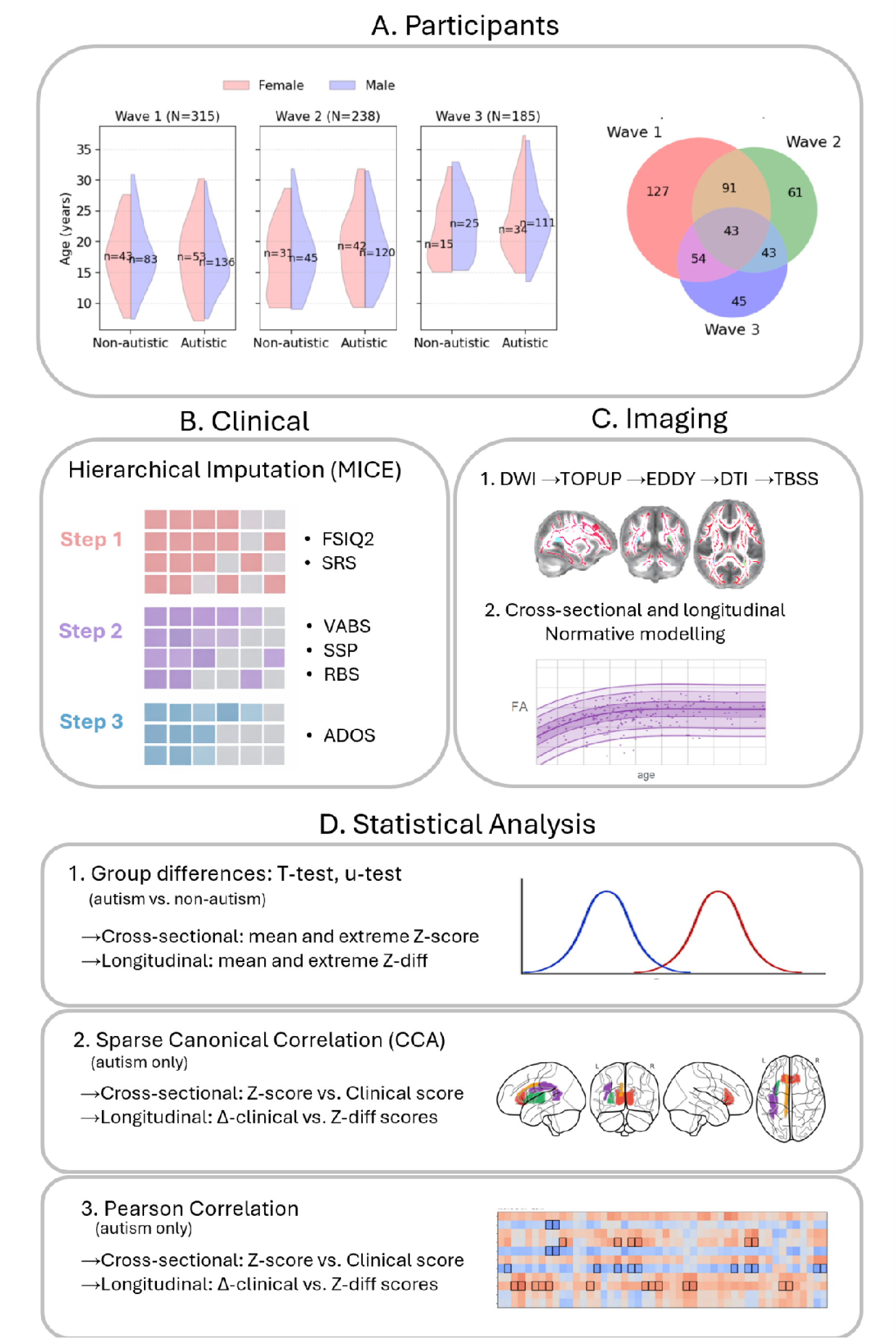
Methods overview pipeline. This study includes data from the three waves of the EU-AIMS LEAP dataset **(A)** spanning the ages 6–37 across five sites. The Venn diagram shows overlap in participant inclusion across waves, indicating the number of participants with available data at one, two, or all three waves. The core clinical data **(B)** has been completed using a customised hierarchical imputation method that accommodates the study design (see methods). The diffusion MRI data from all available datasets were pre-processed **(C)** using the standard FSL pipeline (see methods) and the FA features were extracted using DTIfit and TBSS, obtaining the mean FA value of the skeletonised tract for 48 white-matter tracts according to the JHU white matter atlas. Normative models were applied to each tract to obtain individual FA deviations both cross-sectionally as well as longitudinally (between pairs of visits). **(D)** The statistical analysis comprised three stages for both the cross-sectional as well as the longitudinal data. In the statistical analysis, first, the group comparison tests (t-test and non-parametric u-test) were applied to the FA deviations obtained from the normative models to determine any significant differences between autistic and non-autistic participants. Next, the FA deviations were linked to the core clinical variables in a CCA and Pearson correlation analysis using only autistic participants.

### 2.1. Participants and Study Design

The data were obtained from the EU-AIMS Longitudinal European Autism Project (LEAP), a multicentre programme designed for identifying biological markers of autism across development.^20,21^ Data were collected at five European centres: the Institute of Psychiatry, Psychology and Neuroscience, King’s College London (UK); Radboud University Medical Centre (Nijmegen, the Netherlands); the Autism Research Centre, University of Cambridge (UK); University Medical Centre Utrecht (the Netherlands); and the Central Institute of Mental Health, Medical Faculty Mannheim, University of Heidelberg (Germany). Analyses included 544 individuals (Wave 1: 189 autistic 162 non-autistic; Wave 2: 162 autistic136 non-autistic; Wave 3: 145 autistic 84 non-autistic), aged 7–37 years, with mean intervals of ∼1.5 years (W1–W2) and ∼6 years (W2–W3). The study was approved by local ethics committees at each site, and all participants or guardians provided written informed consent. Demographic distributions are shown in Figure 1A, and demographic information is presented in Supplementary Figure 1 and Supplementary Table 1.

### 2.2. Diffusion MRI

Single-shell diffusion-weighted data (b = 0/1500 s/mm², 60 directions, 2 mm isotropic voxels) were acquired using echo-planar imaging (EPI) sequences. Preprocessing included noise reduction, motion and eddy-current correction, and susceptibility distortion correction using FSL’s eddy.^43–45^ Diffusion tensors were fitted with DTIfit to generate FA maps, which were processed using Tract-Based Spatial Statistics (TBSS).^46^ Each participant’s FA map was projected onto the mean FA skeleton. White matter tracts were segmented using the Johns Hopkins University (JHU) white matter atlas, yielding 48 tracts for which mean FA values were extracted. Full acquisition and preprocessing details are available in Supplementary Text 1.^47^

### 2.3. Normative Modelling

We used normative modelling to quantify individual deviations in white-matter organisation relative to population expectations. For each of the 48 JHU tracts, existing pre-trained lifespan FA models using warped Bayesian Linear Regression were applied to predict expected FA values conditional on age, sex, and site, from which individual Z scores were derived (scores where |Z| > 2.6 were considered extreme deviations).^33,41^ To harmonise model predictions across sites, small wave-specific adaptation sets of non-autistic participants were used to recalibrate normative model intercepts and variance estimates. The adaptation sets consisted of 12 participants per site in each wave, yielding 36 participants for Wave 1, 60 for Wave 2, and 44 for Wave 3, following transfer-learning principles we have reported and evaluated previously (see Supplementary text 2).^48^ Longitudinal deviations were estimated following Bučková et al.^49^ For each tract and wave pair, we computed a Z-diff score to predict how an individual’s change differed from the normative change. Additional quality control (QC) of diffusion data was performed using a normative modelling framework to detect tract-level artefacts and outliers developed in our prior work.^50^ Additional methodological details are described in Supplementary text 2: Normative Modelling.

### 2.4. Clinical Data and Imputation

Clinical and behavioural assessments followed the EU-AIMS LEAP protocol.^20,21^ For this study, we selected measures consistently available across all three waves: These included the Full-Scale Intelligence Quotient II (FSIQ2, an estimated IQ measure harmonised across Wechsler scales), the Autism Diagnostic Observation Schedule Second Edition (ADOS-2), Social Affect calibrated severity score (SA-CSS), the ADOS-2 Restricted and Repetitive Behaviours score (RRB-CSS), and the ADOS-2 Total Calibrated Severity Score (CSS Total), the Social Responsiveness Scale (SRS), the Repetitive Behaviours Scale–Revised (RBS-R), the Short Sensory Profile (SSP), and the Vineland Adaptive Behaviour Scales Second Edition (VABS-II), including the Adaptive Behaviour Composite (ABC), Communication, Daily Living Skills, and Socialisation domains.^51–58^ For longitudinal analyses, clinical and behavioural change scores were calculated between wave pairs. To handle incomplete data, we developed a hierarchical, multi-step imputation approach that accounted for structured missingness based on the Multivariate Imputation by Chained Equations (MICE) software.^59^ For more details about the missingness assessment and imputation procedure see Supplementary text 3.

### 2.5. Statistical Analyses

Analyses were performed separately for cross-sectional (clinical and Z) and pair-wise longitudinal (Δ-clinical and Z-diff) scores across the three LEAP waves/wave-pairs. Group mean differences were tested using two-tailed t-tests, while the proportion of extreme deviations (|Z| > 2.6) per group were compared using Mann–Whitney U-tests for the proportion of extreme deviations, with Benjamini–Hochberg false discovery rate (FDR) correction across tracts (n=48).^49,60^

To examine multivariate brain-behaviour associations, we used canonical correlation analysis (CCA) linking clinical measures and FA deviations for each wave both cross-sectionally as well as longitudinally (between wave pairs).^61^ Model performance was evaluated out-of-sample through ten train–test (70/30) splits, and significance was determined via 1,000 permutations of the clinical data, repeating the entire procedure after permutation. Feature stability was quantified as the proportion of splits in which each variable showed non-zero weight; features selected in ≥70% of splits were considered stable.

To contextualise and assist interpretation of our multivariate findings we conducted univariate correlations between (Δ-)clinical and Z(diff) scores. This was done with each of the ten imputed datasets and combined using Rubin’s Rule, following the implementation described by Marshall et al., with FDR correction applied across tracts.^62,63^ Comprehensive statistical details are provided in Supplementary text 4.

## 3. Results

### 3.1. Normative modelling

The normative models were successfully adapted to the LEAP data and fitted across all 48 white matter tracts, modelling FA trajectories across age (Figure 2A). The model fits varied across waves, with mean explained variance μ_1_= 0.44 in Wave 1, μ_2_= 0.50 in Wave 2, and μ_3_= 0.12 in Wave 3, indicating broadly comparable model fits across waves. The Z score distributions were close to symmetrical (mean skew ≈ 0) but showed moderate positive kurtosis (≈ 0.8–3.8), indicating slightly heavier tails than would be expected under a perfect Gaussian distribution (See Supplementary Figure S1). It is reasonable that we see less explained variance in this analysis and heavier-tailed distributions due to the narrow age range, conservative adaptation set, partial site overlap, and the high variability typical of paediatric diffusion acquisitions relative to our entire lifespan cohorts.^41^

**Figure 2.**
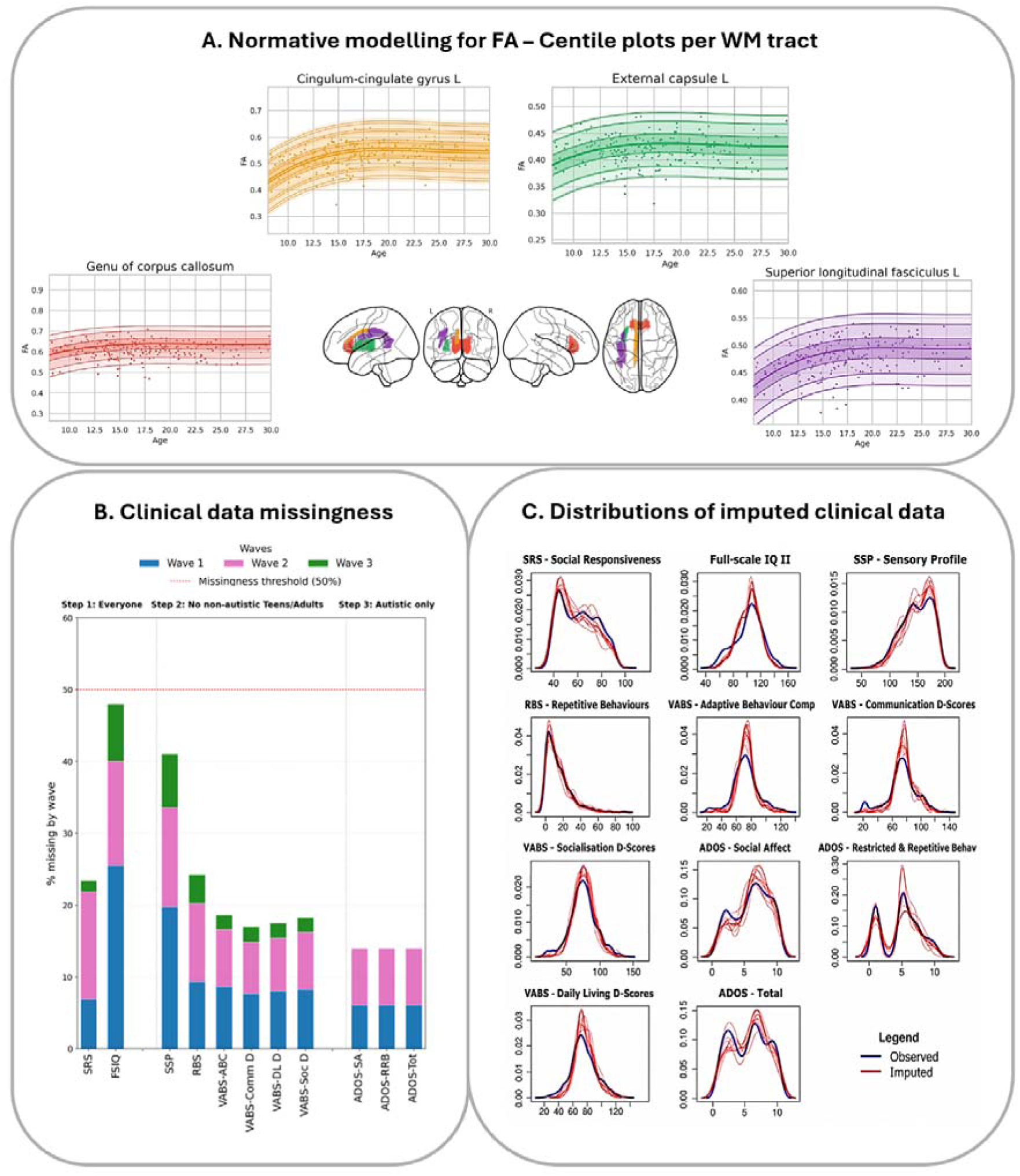
Normative modelling and clinical data completeness. **(A)** *FA normative models*. Centile plots from select representative white matter tracts (frequently occurring in the CCA and correlation results) show a good fit with minimal outliers. The lifespan trend shows the expected white matter maturation curve with rapid increase in childhood followed by stabilisation in adulthood. **(B)** *Clinical data missingness.* The core clinical traits we investigated showed moderate missingness levels especially in Wave 1 and 2 where the number of participants was larger. Wave 3 has a relatively more complete profile with no missing values within the ADOS cluster. **(C)** *Imputed clinical data distributions.* To mitigate the missingness problem and increase the usability of the dataset, we imputed the data longitudinally, obtaining estimates (red curves) which closely follow the distributions of the observed values (blue curve).

### 3.2. Group Comparisons

Across cross-sectional and longitudinal analyses, no tract-wise group differences survived correction for multiple comparisons. Nominal effects were observed in a small number of tracts, including the cerebral peduncles, external capsule, cingulum-hippocampus, corpus callosum, internal capsule, posterior corona radiata, and tapetum, but these did not remain significant after FDR correction. Full uncorrected results are reported in Supplementary Table 2.

### 3.3. Canonical correlation analysis (CCA)

Despite relatively high proportions of missing data (Figure 2B), our hierarchical imputation strategy provided high-quality imputations that respect the original data distribution (Figure 2C). Cross-sectional CCA results are summarised in Figure 3 and reveal significant out-of-sample correlations between FA deviations (48 tracts) and the imputed clinical measures (described in the Methods section) for Waves 1 and 2, within the autism group, but not Wave 3. In Wave 1, the first component shows an out-of-sample canonical correlation of r = 0.26, consistently selecting FA loadings in the cingulum L, anterior limb of the internal capsule R, external capsule L, and genu of the corpus callosum for the first component. Wave 2 showed a similar pattern (r = 0.32), consistently selecting the bilateral cingulum, superior longitudinal fasciculus, splenium of the corpus callosum, and cerebral peduncles for the first component.

**Figure 3.**
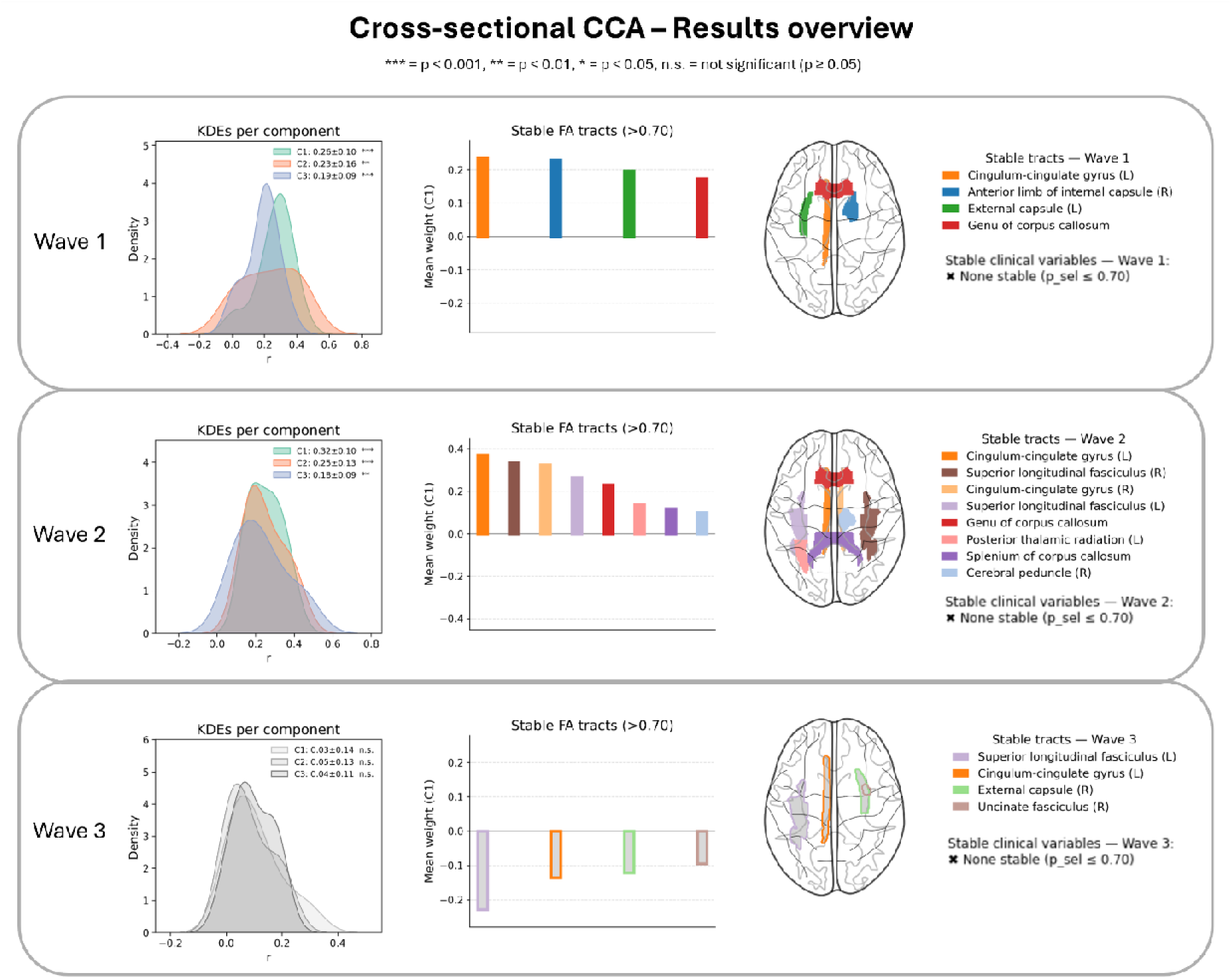
Cross-sectional CCA results overview. The sparse CCA using the cross-sectional data shows recurrent associations linking individual FA deviations to core clinical traits in autism. These associations are wave dependent with Wave 1 and 2 showing significant results under permutation testing (left), whereas effects are attenuated in Wave 3. The stability of the implicated features also varies in magnitude and direction with the imaging features dominating the multivariate associations (centre and right), suggesting that clinical features have a general contribution rather than being single dominant drivers. Across waves, stable contributions consistently involve long-range association and commissural pathways, including the cingulum bundle, superior longitudinal fasciculus, external capsule, and corpus callosum.

**Figure 4.**
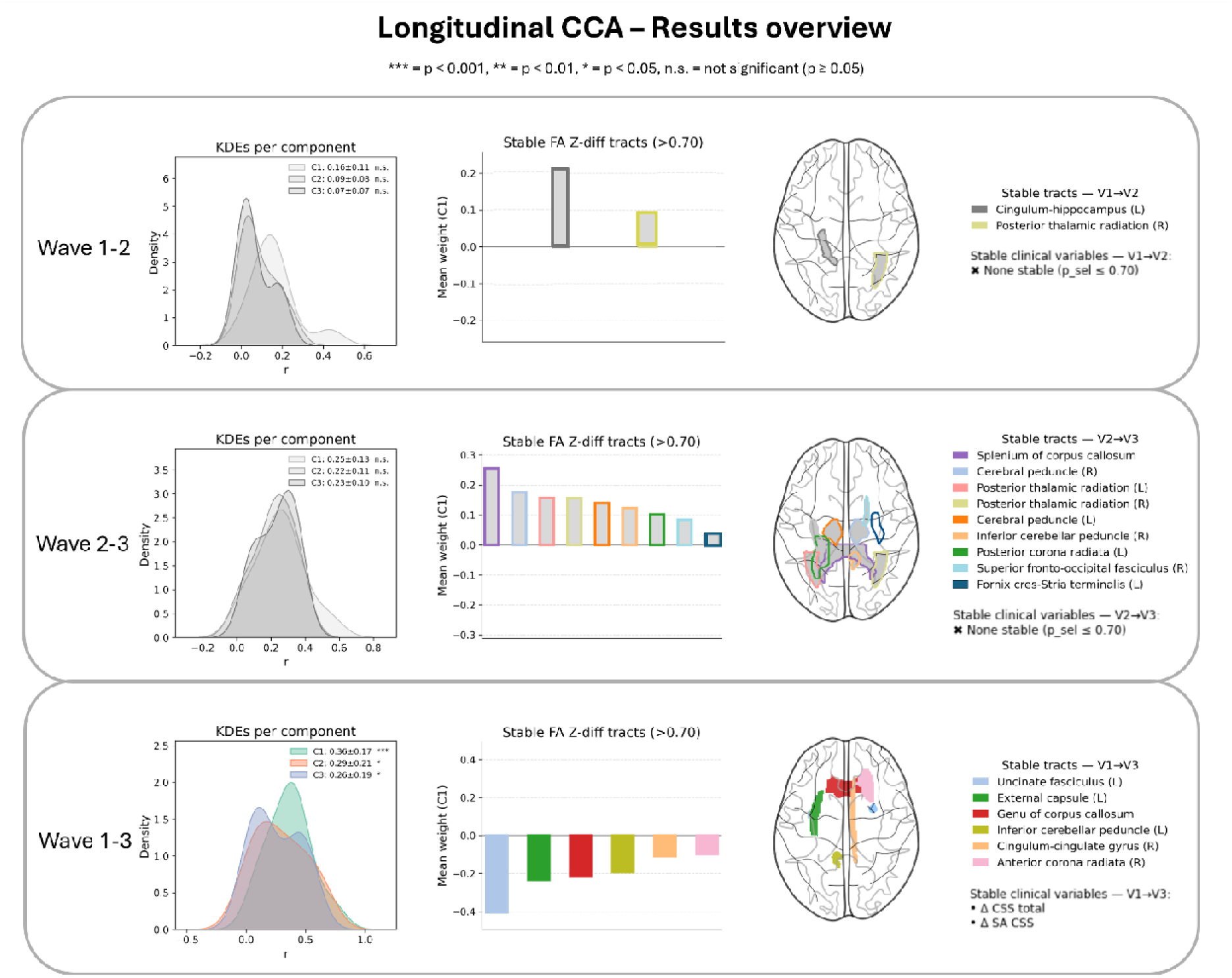
Longitudinal CCA results overview. Sparse CCA of the longitudinal data reveals fewer brain–behaviour associations than the cross-sectional analysis. Only one wave pair (Wave 1–3) was significant under permutation testing (left), suggesting that longitudinal brain–behaviour relationships may vary across developmental intervals. Even though the signal is attenuated, tract-level features were selected across wave pairs, involving association, commissural, and projection fibres (centre and right). In Wave 1–3, changes in ADOS total severity and social-affect severity were consistently selected, indicating that the significant longitudinal component was primarily linked to change in autism trait severity.

Across the longitudinal analyses, only the Wave 1-3 comparison had significant canonical correlations. The first component showed a moderate association (r = 0.36) and included stable FA Z-diff loadings in several projection and association tracts, such as the genu of the corpus callosum, left inferior cerebellar peduncle, bilateral anterior corona radiata, right sagittal stratum, bilateral external capsule, left uncinate fasciculus, and right tapetum. This was also the only analysis in which clinical variables were stably selected, specifically ΔCSS Total and ΔSA-CSS.

### 3.4. Pooled Pearson Correlation Analysis

The cross-sectional analyses showed significant associations between specific tract Z scores and multiple clinical dimensions across waves (Figure 5, left panel). In Wave 1, SSP and ADOS-RRB showed widespread negative associations, while VABS exhibited positive correlations, particularly in cerebellar and fronto-limbic tracts. Several of these effects survived FDR correction, including in the uncinate fasciculus, cerebellar peduncles, and cingulum. Wave 2 showed widespread correlations between VABS and many of the same tracts as in Wave 1 but also parts of the internal capsule, thalamic radiation and sagittal stratum. In Wave 3, associations persisted but were weaker and less robust after correction.

**Figure 5.**
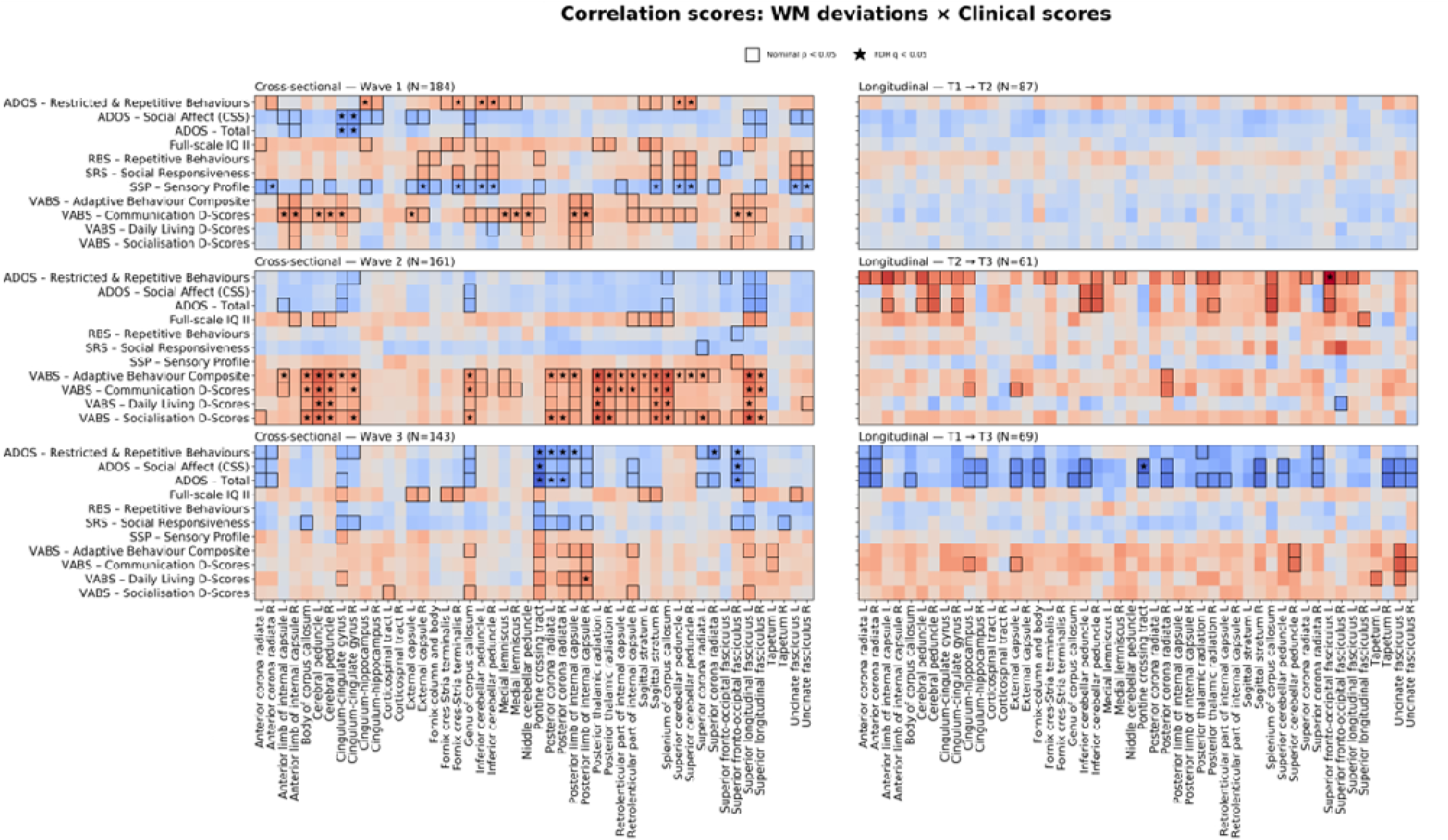
Pearson correlations. These figures present heatmaps of the correlation scores between the FA deviations (Z score or Z-diff) for each of the 48 tracts and each core clinical variable, in autistic participants. The bold frames around a cell represent a nominally significant score, while the star marks the scores significant after FDR testing. The left panel reveals stronger overall associations in the cross-sectional analyses compared to the longitudinal results (right panel) including both broader regions of correlations as well as more significant relationships surviving FDR correction. In the cross-sectional analysis: across tracts, the VABS domains show a predominantly positive correlation pattern with association, commissural and projection fibre tracts, whilst ADOS and SRS/RBS reveal mixed and wave-dependent scores and directions.

Longitudinally, we observed less pronounced effects, with only a few nominal associations emerging between changes in clinical scores and Z-diff scores (Figure 5, right). Wave 2-3 comparisons showed nominal correlations in ΔADOS scores across cingulate and projection pathways, with one tract (left superior fronto-occipital fasciculus) surviving FDR correction. Moderate ΔADOS correlation patterns were also seen in Wave 1-3 for fronto-limbic and thalamic pathways. The pontine crossing tract emerged as a key region for longitudinal change.

## 4. Discussion

In this study, we used normative modelling to characterise the white-matter organisation in autism and its relationship to continuous behavioural traits, both cross-sectionally and longitudinally. We showed that in autism, individual deviations, rather than group-mean differences, accounted for variance in clinical traits within fronto-limbic, callosal, and cerebellar–thalamo-cortical pathways, implicating adaptive, sensory, and behavioural dimensions.

The cross-sectional analysis revealed relatively consistent patterns: the CCA results showed stable component loadings in the cingulum, corpus callosum, and cerebellar tracts across Waves 1 and 2, while the pooled Pearson correlations pointed to direction-specific associations between FA deviations and sensory (SSP), adaptive (VABS), and behavioural (ADOS) traits. VABS correlated with tracts such as the cingulum, corpus callosum, internal capsule and cerebellar peduncles. However, ADOS displayed both mixed and recurrent associations, positive for restricted behaviours and negative for social domains, in the fronto-limbic and cerebellar regions. The direction of clinical score scaling should be considered when interpreting these associations. Higher VABS and SSP scores indicate better adaptive and sensory functioning, whereas higher ADOS scores indicate greater autism symptom severity. Thus, higher FA was associated with better adaptive functioning but greater sensory difficulties, suggesting domain-specific rather than a uniform interpretation of higher FA, consistent with existing research.^65^

Although similar patterns were observed across the three waves, the statistical significance varies, especially in Wave 3, as many of the nominally significant findings did not survive correction for multiple comparisons. This is likely an effect of sample size variation and increased developmental heterogeneity, as Wave 3 no longer spanned childhood and consisted exclusively of adolescents and adults. In particular, adolescence is characterised by non-linear white-matter maturation and increased inter-individual variability, which may reduce the sensitivity to detect linear brain–behaviour associations.^41,66,67^ The attenuation of effect sizes observed in Wave 3 is therefore more plausibly interpreted as reflecting variation in developmental timing rather than loss of biological signal.

Many of our cross-sectional results are supported by existing research. For example, the relationship between autism trait severity and alterations in limbic and emotion-related tracts reported by Gibbard et al. is in line with our CCA findings in the cingulum (all three waves) and uncinate fasciculus (at Wave 3).^68^ The correlation between sensory processing and FA in association and limbic tracts described by Pryweller et al. also matches the negative sensory correlations we observed in Wave 1 in the pooled Pearson correlation analysis.^69^ Recently, several other neuroimaging studies have focused on individualised models in autism. Using normative modelling, Zabihi et al. demonstrated that the brains of autistic individuals are characterised by individual deviation in cortical thickness and revealed clinically relevant subtypes, while group-level differences proved to be minimal.^37,70^ At the white-matter level, Aoki et al. showed dimensional clinical measures can reveal white-matter patterns beyond those captured by diagnostic comparisons alone, particularly in the corpus callosum.^42^

Longitudinally, our analysis indicated that in autism white-matter microstructural differences and clinical traits change together. In the CCA analysis, the Wave 1–3 pair showed that FA changes in tracts including the inferior cerebellar peduncle, uncinate fasciculus, genu of the corpus callosum, external capsule and anterior corona radiata were associated with a change in ADOS, providing evidence that these pathways represent an ongoing relationship between evolving brain structure and autistic traits over time. ADOS also emerged as nominally significant in the pooled Pearson correlation analysis as well, showing associations with some of the previously encountered tracts including the anterior corona radiata, external capsule (left), inferior cerebellar peduncle, uncinate fasciculus and genu of the corpus callosum in the same Wave 1-3 pair. Compared to the cross-sectional analyses, our longitudinal analyses showed fewer significant associations, especially after correcting for multiple comparisons, reflecting reduced statistical power due to smaller sample sizes, partial overlap across waves.

While longitudinal studies on white-matter organisation in autism are scarce, two relevant articles emerge with important findings. First, Zhang et al. have demonstrated that lower FA in the superior longitudinal fasciculus is associated with increased communication difficulty and that FA values in the superior thalamic radiation are good predictors of diagnostic outcome.^30^ Similarly, our findings show that higher VABS scores were linked to stronger FA in association and thalamo-cortical tracts. Second, Andrews et al. also demonstrated that in children aged 2.5–7, slower FA development in the cingulum, superior longitudinal fasciculus, internal capsule and splenium correlated with changes in ADOS severity scores, supporting the brain-behaviour relationships observed in our longitudinal analysis.^29^

A few technical considerations and limitations must be mentioned. First, FA is a non-specific measure reflecting multiple microstructural properties, including myelination, axonal coherence, and fibre organisation, and deviations from normative FA trajectories are better understood as generally reflective of white matter organisation.^31,66^ Furthermore, sample size and data completeness pose an important limitation. At the same time, greater overlap across waves and reduced clinical data missingness would improve longitudinal power and interpretability. Although multiple imputation go some way to addressing this, they do not replace truly observed data. Nevertheless, this approach offers a robust and principled way to estimate missing values while preserving sample size and maintaining statistical power.

In conclusion, by placing individual FA values against developmentally informed reference trajectories, normative modelling makes it possible to interpret white-matter variation as clinically meaningful individual deviation both cross-sectionally and longitudinally. In autism, these deviations mapped onto adaptive, sensory, and behavioural variation, supporting a dimensional view of white-matter organisation that is sensitive to heterogeneity across development.

## Abbreviations

ABC: Adaptive Behaviour Composite
ADOS: Autism Diagnostic Observation Schedule
B-spline: Basis spline
BLR: Bayesian Linear Regression
CCA: Canonical Correlation Analysis
CSS Total: Calibrated Severity Score (ADOS Total)
DSM-5: Diagnostic and Statistical Manual of Mental Disorders, Fifth Edition
DTI: Diffusion Tensor Imaging
DTIFIT: Diffusion Tensor Imaging Fitting tool (FSL)
DWI: Diffusion-Weighted Imaging
FA: Fractional Anisotropy
FDR: False Discovery Rate
FSIQ2: Full-Scale Intelligence Quotient, two-subtest form
FSL: FMRIB Software Library
LEAP: Longitudinal European Autism Project
MICE: Multivariate Imputation by Chained Equations
MRI: Magnetic Resonance Imaging
msCCA: Multi-view sparse Canonical Correlation Analysis
QC: Quality Control
RBS: Repetitive Behaviours Scale
RRB-CSS: Restricted and Repetitive Behaviours Calibrated Severity Score
SA-CSS: Social Affect Calibrated Severity Score
SRS: Social Responsiveness Scale
SSP: Short Sensory Profile
TBSS: Tract-Based Spatial Statistics
VABS: Vineland Adaptive Behaviour Scales
Z: Normative deviation score
Z-diff: Normative deviation of longitudinal change

## Funding

This research was supported by grants from the European Research Council (ERC, grant “MENTALPRECISION ”10100118) and NIH grant number 1R01MH130362-01A1.

The results leading to this publication have received funding from the Innovative Medicines Initiative 2 Joint Undertaking under grant agreement No 115300 and No 777394 for the projects EU-AIMS and AIMS-2-TRIALS. This Joint Undertaking receives support from the European Union’s Horizon 2020 research and innovation programme and EFPIA and AUTISM SPEAKS, Autistica, SFARI. Any views expressed are those of the authors and not necessarily those of the funders (IHI-JU2).

This work was supported by EU-AIMS (European Autism Interventions), which received support from the Innovative Medicines Initiative Joint Undertaking under grant agreement no. 115300, the resources of which are composed of financial contributions from the European Union ’ s Seventh Framework Programme (grant FP7/2007-2013), from the European Federation of Pharmaceutical Industries and Associations companies ’ in-kind contributions, and from Autism Speaks.

## Author contributions

Conceptualisation: RC, NF, AM, CB

Methodology: RC, NF, AM, BRB, CB

Validation: RC, NF, AM, BRB, CB

Software: RC

Formal analysis: RC, NF, AM, BRB, CB

Visualisation: RC

Supervision: NF, AM, CB

Investigation: RC, NF, AM, BRB, CB, FD, TC, EL, BO, LCM, IY, RS, SD, TB, SBC, DGMM, JKB, EU-AIMS LEAP Group

Data curation: FD, EL, BO, LCM, IY, RS, SD, TB, SBC, DGMM, JKB

Writing—original draft: RC

Writing—review & editing: RC, NF, BRB, FD, TC, EL, BO, LCM, IY, RS, SD, TB, SBC, DGMM, JKB, CB, AM

## Financial Disclosure or Competing interests

CB is director and shareholder for SBGneuro. TC has received consultancy fees from F. Hoffmann-La Roche Ltd. and royalties from Sage Publishing and Guilford Press.

None of the other authors report any biomedical financial interest or potential conflicts of interest.

## Data sharing statement

LEAP data are accessible through the EU-AIMS consortium; external access requires submission of an approved analysis proposal under the consortium’s data-sharing policies. Fractional anisotropy (FA) normative models used in this study are publicly available via https://pcnportal.dccn.nl/ for reuse by the scientific community. All analysis code and normative modelling pipelines developed in this work are openly available on GitHub at https://github.com/ramonacirstian.

## Ethics and privacy

All datasets included in this thesis were collected under the approval of the relevant local ethics committees of the originating studies, and all participants provided informed consent for use and sharing and reuse of their data in accordance with the Declaration of Helsinki. The work presented here involves secondary analysis of existing, de-identified data (no data was collected for the purpose of this dissertation) and therefore did not require additional ethical approval. The privacy of all participants was ensured through randomised unidentifiable subject codes

## Use of AI in Publication and Research

Artificial intelligence tools (ChatGPT, GPT-5, OpenAI) were used for limited assistance with programming troubleshooting and language editing. The study conception, design, analyses, interpretation, and manuscript drafting were conducted by the authors. AI tools did not generate data, results, or scientific conclusions. The authors affirm that the work is original and accept full responsibility for its content.

## Notes

### Competing Interest Statement

CB is director and shareholder for SBGneuro. None of the other authors report any biomedical financial interest or potential conflicts of interest.

### Summary of Updates

Figure 1 was added, Supplementary Table 1 added

