## supplement for "Linking White-Matter Development to Clinical Variation in Autism: Longitudinal Normative Modelling of Fractional Anisotropy (FA) in the EU-AIMS LEAP Cohort"

Ramona Cirstian* et al.

*Corresponding author.

### Supplementary Table T1: Demographic and Clinical Information

| Wave 1 | | |
| --- | --- | --- |
| Measure - Mean (SD) | Autistic | Non-autistic |
| Participants | N = 189 (F = 53, M = 136) | N = 162 (F = 61, M = 101) |
| Age | 17.23 (5.21) | 17.45 (5.25) |
| SRS – Social Responsiveness | 71.08 (12.06) | 48.21 (8.38) |
| SSP – Sensory Profile | 140.20 (26.74) | 177.47 (14.87) |
| RBS – Repetitive Behaviours | 15.85 (13.50) | 2.40 (4.67) |
| VABS – Adaptive Behaviour Composite | 70.05 (14.77) | 85.45 (25.61) |
| VABS – Communication D-Scores | 75.72 (17.76) | 85.23 (27.78) |
| VABS – Daily Living D-Scores | 71.80 (16.61) | 85.21 (22.59) |
| VABS – Socialisation D-Scores | 68.59 (16.30) | 88.42 (24.24) |
| ADOS – Social Affect | 6.04 (2.70) |  |
| ADOS – Restricted & Repetitive Behaviours | 4.86 (2.79) |  |
| ADOS – Total | 5.44 (2.83) |  |
| Full-scale IQ II | 97.63 (22.13) | 104.19 (20.70) |
| Wave 2 | | |
| Measure - Mean (SD) | Autistic | Non-autistic |
| Participants | N = 162 (F = 42, M = 120) | N = 136 (F = 50, M = 86) |
| Age | 18.88 (5.69) | 17.59 (5.77) |
| SRS – Social Responsiveness | 70.58 (11.38) | 45.07 (6.61) |
| SSP – Sensory Profile | 146.10 (25.05) | 179.78 (9.86) |
| RBS – Repetitive Behaviours | 13.43 (13.77) | 1.24 (2.40) |
| VABS – Adaptive Behaviour Composite | 73.41 (15.69) | 100.40 (30.24) |
| VABS – Communication D-Scores | 75.37 (18.99) | 93.73 (38.43) |
| VABS – Daily Living D-Scores | 76.24 (16.58) | 91.82 (28.59) |
| VABS – Socialisation D-Scores | 75.65 (17.01) | 103.50 (30.71) |
| ADOS – Social Affect | 5.89 (2.75) |  |
| ADOS – Restricted & Repetitive Behaviours | 5.09 (2.65) |  |
| ADOS – Total | 5.35 (2.88) |  |
| Full-scale IQ II | 99.82 (19.76) | 113.31 (16.12) |
| Wave 3 | | |
| Measure - Mean (SD) | Autistic | Non-autistic |
| Participants | N = 145 (F = 34, M = 111) | N = 84 (F = 28, M = 56) |
| Age | 23.60 (5.45) | 22.41 (5.94) |
| SRS – Social Responsiveness | 64.24 (12.10) | 45.38 (6.71) |
| SSP – Sensory Profile | 150.24 (28.01) | 182.36 (9.61) |
| RBS – Repetitive Behaviours | 14.59 (12.99) | 2.23 (5.81) |
| VABS – Adaptive Behaviour Composite | 68.65 (15.90) | 41.00 (16.75) |
| VABS – Communication D-Scores | 67.24 (22.83) | 33.17 (10.68) |
| VABS – Daily Living D-Scores | 70.80 (17.64) | 48.00 (19.42) |
| VABS – Socialisation D-Scores | 75.08 (16.97) | 47.50 (21.03) |
| ADOS – Social Affect | 5.78 (3.00) | 2.62 (2.38) |
| ADOS – Restricted & Repetitive Behaviours | 5.31 (2.98) | 2.65 (2.37) |
| ADOS – Total | 5.42 (3.12) | 2.27 (2.16) |
| Full-scale IQ II | 102.22 (20.25) | 107.55 (21.09) |

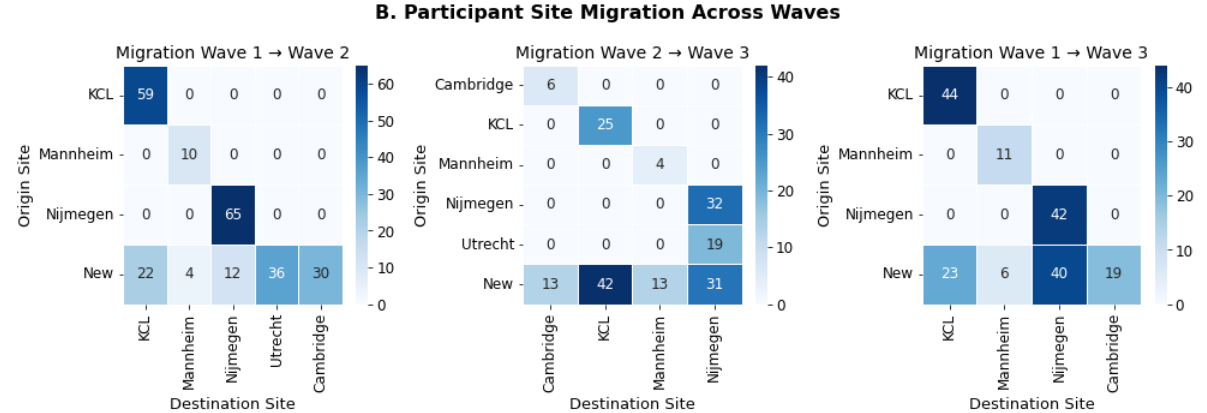

### Supplementary Figure S1: Participation across waves and sites

**Supplementary Figure 1. Site migration between waves**. Heatmaps depict origin (rows) → destination (columns) site transitions for participants observed at both waves (W1→W2, W2→W3, and W1→W3). The New row counts subjects present at the later wave but absent at the preceding wave, allocated by their destination site.

Data were collected at five European centres:

- The Institute of Psychiatry, Psychology and Neuroscience, King’s College London (KCL, UK)
- Radboud University Medical Centre (Nijmegen, the Netherlands)
- The Autism Research Centre, University of Cambridge (UK)
- University Medical Centre Utrecht (the Netherlands)
- The Central Institute of Mental Health, Medical Faculty Mannheim, University of Heidelberg (Germany)

### Supplementary text 1: Diffusion MRI

Single-shell diffusion-weighted images were acquired across all participating sites using a spin-echo echo-planar imaging (EPI) sequence with the following parameters (site range): repetition time (TR) = 1200 ms; echo time (TE) = 65–103 ms; flip angle = 90°; 72 slices; voxel size = 2 × 2 × 2 mm³; and b-values = 0 and 1500 s/mm² with 60 diffusion directions and 6 b₀ volumes.

All data underwent a harmonised preprocessing and quality control protocol as previously reported by Mei et al. (1). Noise removal was performed using the Marchenko–Pastur principal component analysis (MP-PCA) method (2), followed by Gibbs ringing correction (3). Eddy-current distortions, motion, and slice outliers were then corrected with FSL’s eddy (4), including slice-to-volume motion correction (5).

To address site-specific signal issues, datasets from Nijmegen and Mannheim were corrected using an in-house implementation of Koay et al. (6). Remaining slice-wise signal dropouts in b₀ images were identified using an interquartile range (IQR) outlier threshold (Q3 + 1.5×IQR) and replaced with the corresponding slice median. To further harmonise diffusion signal profiles across sites, the data were resampled using a real symmetric spherical harmonics (SH) representation limited to l = 6, which reduced high-frequency noise and improved cross-site consistency. For Siemens scanners showing residual Gibbs ringing, a final b₀ correction was applied using ExploreDTI (7). Data quality was assessed using visual inspection and quantitative metrics following Bastiani et al.(8) . Participants with head motion exceeding 4 mm translation or more than 6% outlier slices were excluded.

Diffusion tensor models were fitted using FSL DTIFIT to derive voxelwise fractional anisotropy (FA) maps. Tract-Based Spatial Statistics (TBSS) (9) were then applied, including registration to standard space, projection onto a mean FA skeleton, and generation of individual skeletonised FA maps.

White matter segmentation was performed using the Johns Hopkins University (JHU) white matter atlas (10), delineating 48 tracts. Mean FA values were extracted along the skeleton of each tract. Diffusion MRI data quality was further evaluated through tract-wise normative modelling of FA values, with extreme outliers (|Z| > 4) subjected to visual inspection.

### Supplementary text 2: Normative Modelling

##### FA Normative Models

To obtain the individual variations in white matter organisation we used FA normative models previously developed, trained on 12457 participants across the lifespan (0-100 years) with covariates including sex, age and site (11). Separate models were trained on 48 white matter tracts (using the JHU atlas (10)) using warped Bayesian Linear Regression with age modelled using a 3rd-order B-spline basis as described by Fraza et al. (12). We applied a Sinh–Arcsinh warping transformation to account for non-linearity (equation 1) while site effects were modelled as fixed effects which allowed us to estimate site-specific variances. Each model predicted tract-specific means and standard deviations, from which individual deviation (Z) scores were computed (equation 2):

$\varphi_{SinhArcsinh}\left( \boldsymbol{y};\boldsymbol{\gamma} \right)=\sinh\left( b*arcshinh\left( \boldsymbol{y} \right)+\varepsilon*b \right)$	(equation 1)

$z_{nd}=\frac{y_{nd}-ŷ_{nd}}{\sqrt{\sigma_{d}^{2}+{(\sigma_{*}^{2})}_{d}}}$	(equation 2)

In equation 1: $a=-\varepsilon*b$ and $\varphi_{Affine}\left( \boldsymbol{y};\boldsymbol{\gamma} \right)=a+b\boldsymbol{y}$.

In equation 2: *n* denotes each subject while *d* denotes each white matter tract and $ŷ_{nd}$ is the predicted mean while $y_{nd}$ denotes the true response. The estimated noise variance (i.e. reflecting variation in the data) is denoted by $\sigma_{d}^{2}$ and the variance attributable to modelling uncertainty for the 𝑑-th voxel is denoted by ${(\sigma_{*}^{2})}_{d}$.

##### Application to LEAP Data

We applied the FA normative models to the LEAP dataset to compare the de FA deviation in LEAP participants against the larger reference cohort previously modelled. To harmonise model predictions across sites, small wave-specific adaptation sets of control participants were used to recalibrate normative model intercepts and variance estimates, following transfer-learning principles we have reported and evaluated previously (13). For each wave, the site-matched adaptation sets were assembled from typically developing controls (12 subjects per site), wave unique controls were prioritised while controls within multiple waves were only added when the required number (12) was not met; all remaining participants (autistic and non-adaptation controls) comprised the test sets. The adaptation and test set sizes were as follows:

- Wave 1:
  - Total participants: 351
  - Adaptation set: 36
  - Final test sample: 315
- Wave 2:
  - Total participants: 298
  - Adaptation set: 60
  - Final test sample: 238
- Wave 3:
  - Total participants: 229
  - Adaptation set: 44
  - Final test sample: 185
- Wave 1 & Wave 2: 134 shared participants
- Wave 2 & Wave 3: 86 shared participants
- Wave 1 & Wave 3: 97 shared participants

##### Longitudinal Normative Modelling

Longitudinal deviations were estimated following Bučková et al.(14). For each tract and wave pair, we computed a Z-diff score to predict how an individual’s change differed from the normative change according to the following equation:

$$Z_{diff}=\frac{\left( Y_{v2}-Y_{v1} \right)-\left( \hat{Y_{v2}}-\hat{Y_{v1}} \right)}{\sigma_{\Delta\varepsilon}}$$

Where, $Y_{v1}$ and $Y_{v2}$ are the observed FA values at visits 1 and 2, and $\hat{Y_{v1}}$ and $\hat{Y_{v2}}$ are the corresponding normative predictions, while the denominator $\sigma_{\Delta\varepsilon}$ denotes the standard deviation of residual change among controls.

##### Model Evaluation and Quality Control

Model performance was evaluated following the framework outlined in Cristian et al. (15), using metrics of explained variance, skewness, and kurtosis to assess model fit across tracts which are depicted in Supplementary Figure S2 - Normative Modelling Fit Metrics Summary. Automated quality-control procedures were used to flag potential outlier Z-scores, which were then visually inspected for plausibility. One participant with a scanner artefacts was excluded from Wave 1 and the rest of the data was retained. For visualisation, we generated centile plots showing tract-wise normative ranges and individual deviations for each wave. All preprocessing and model adaptation was implemented in Python using the PCN toolkit.

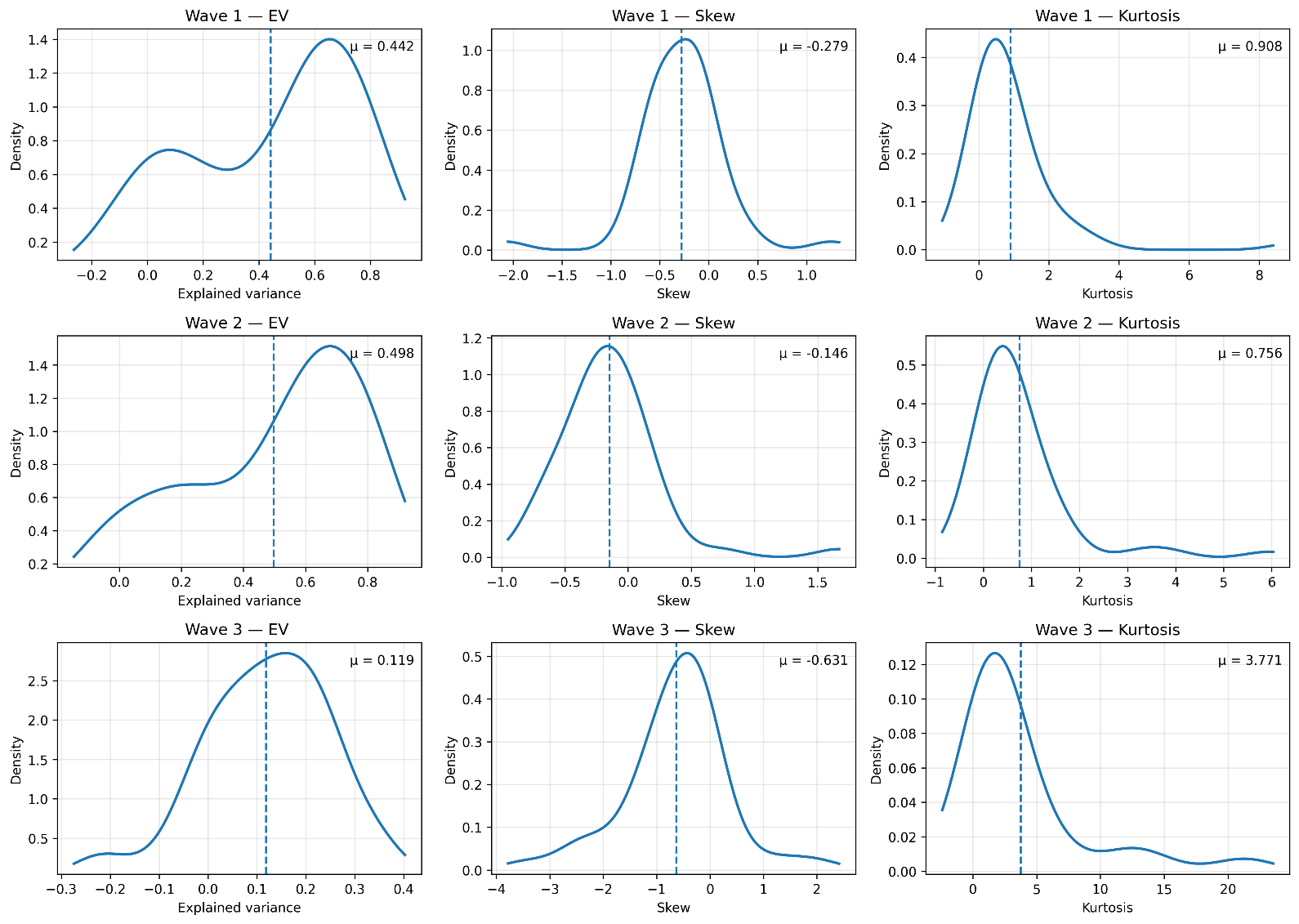

### Supplementary Figure S2 - Normative Modelling Fit Metrics Summary

**Supplementary Figure S1. Normative Modelling Fit Metrics Summary.**

Distributions of tract-wise model calibration metrics for Waves 1-3. Columns show the explained variance (EV), skewness, and kurtosis of tract-level Z-score distributions. The models demonstrated stable calibration across tracts, with mean explained variance μ_1_= 0.44 in Wave 1, μ_2_= 0.50 in Wave 2, and μ_3_= 0.12 in Wave 3. Z-distributions were approximately symmetric (mean skew near 0) but exhibited moderate positive kurtosis (μ ≈ 0.8–3.8), indicating slightly heavier tails than expected under a perfect Gaussian fit.

### Supplementary text 3: Missingness and Imputation

Preparing the LEAP clinical dataset required addressing a complex pattern of missingness that was both random and structured by design. To prepare the data for imputation, we classified participants into ten schedules combining age group (child, adolescent, adult), diagnostic, and intellectual-disability according to the study protocol described by Charman et al. (2017) (16) and the imputation framework of Llera et al. (2022) (17). This label-based structure made it clear which measures were expected for each subgroup and that imputation had to account for systematic absences (not treat them as random gaps). We ensured that no subjects were missing more than 50% of the data by excluding those who exceeded this threshold. Furthermore, none of the variables presented more than 50% missingness across the total number of subjects.

The rationale for our approach was therefore both methodological and conceptual. From previous works on clinical data imputation (17) (18), we learned that attempting to impute all variables in a single model would risk propagating artificial correlations between measures never collected. Therefore, we implemented a hierarchical imputation design that mirrors the LEAP study design.

All imputations were performed in R using the mice package (19) with Classification and Regression Trees (CART) as the predictive method. CART was chosen for its flexibility in capturing non-linear and mixed-type relationships. To accommodate our longitudinal framework, we added subject fixed effects so that each participant’s within-person pattern across waves was preserved during estimation.

The process followed three sequential steps.

- Step 1 imputed cognitive and trait measures (IQ and SRS), including all participants (with complete demographic information).
- Step 2 expanded the model to adaptive, sensory, and behavioural variables (Vineland, -SSP, RBS), for schedules in which these tests were administered (everyone except adult and adolescents TD).
- Step 3 addressed ADOS Calibrated Severity Scores, restricted to the ASD schedules.

Each step was run 10 times (m = 10 imputations per step). From each batch, one completed dataset was chosen at random and passed to the next step. The entire 3-step chain was then repeated 10 independent times, resulting in 10 final fully imputed datasets per step.

### Supplementary text 4: Statistical Analyses

All analyses were performed separately for cross-sectional (Z-scores) and longitudinal (Z-diff) data across the three LEAP waves, resulting in six analytical sets in total.

##### White-Matter Extreme Deviation Analysis

Individual white-matter deviation (Z) and longitudinal change (Z-diff) scores were examined to identify participants with atypical white-matter organisation. Extreme deviations were defined as |Z| > 2.6, corresponding to approximately p ≈ 0.01 under a standard normal distribution. For each tract, we compared the proportion of participants exceeding this threshold between autistic and control groups. Mean tract-wise Z-scores were compared using two-tailed Welch’s t-tests, and group differences in distributional ranks were assessed using Mann–Whitney U tests. In both cases, p-values were corrected for multiple comparisons using the Benjamini–Hochberg false-discovery rate (FDR).

##### Multivariate FA-Clinical Associations (msCCA)

To assess multivariate associations between white-matter microstructure and clinical measures, we employed a multi-view canonical correlation analysis (msCCA) following the framework described by Ing et al. (20). This method identifies sparse linear combinations of variables that maximise shared variance between datasets:

$$max_{w_{1},,w_{2}};w_{1}^{T}X_{1}^{T}X_{2},w_{2}$$

subject to the constraints:

$$\left| w_{1} \right|_{2}=\left| w_{2} \right|_{2}=1,\quad\left| w_{1} \right|_{1}\leq c_{1},;\left| w_{2} \right|_{1}\leq c_{2}$$

Here, $X_{1}$ and $X_{2}$denote matrices of standardised clinical and FA-deviation variables, and the regularisation parameters c_v control sparsity. Equal sparsity penalties were applied to both data views (L1 = [0.5, 0.5]). Three canonical components were extracted (rank = 3).

Model stability was assessed using ten random 70/30 train–test splits, each providing out-of-sample canonical correlations. Feature stability was quantified as the proportion of splits in which each variable exhibited non-zero weight and features selected in ≥70% of splits were considered stable. To test statistical significance, we performed 1000 permutation tests where the clinical data was randomly permuted to destroy cross-view correspondence. Empirical p-values were obtained as the proportion of permuted test correlations greater than or equal to the observed value.

##### Correlation Analysis with Multiple Imputation (Rubin’s Rule)

Correlations between clinical measures and tract-wise FA deviations (or Z-diff values) were computed separately for each imputed dataset and then combined using Rubin’s Rule (21), following the implementation described by Marshall et al. (22), ensuring that both within- and between-imputation uncertainty were reflected in the final estimates.

For each tract–clinical pair i, Pearson’s correlation $r_{i}$ and sample size $n_{i}$ were computed and transformed to Fisher’s *z*:

$$z_{i}=\tanh^{-1} \left( r_{i} \right),\quad U_{i}=\frac{1}{n_{i}-3}$$

where zᵢ is the Fisher-transformed correlation and Uᵢ represents its within-imputation variance.

The pooled estimate was:

$$\bar{z}=\frac{1}{m}\sum_{i=1}^{m} z_{i},$$

$$\bar{U}=\frac{1}{m}\sum_{i=1}^{m} U_{i},$$

$$B=\frac{1}{m-1}\sum_{i=1}^{m} \left( z_{i}-\bar{z} \right)^{2}$$

where z̄ is the mean Fisher’s z across imputations, Ū is the average within-imputation variance, and B is the between-imputation variance reflecting imputation uncertainty.

The total variance and standard error were then:

$$T=\bar{U}+\left( 1+\frac{1}{m} \right)B,\quad SE=\sqrt{T}$$

The test statistic ($Z)$, p-value ($p)$, and pooled correlation ($r_{\text{pooled}})$ were derived as:

$Z=\frac{\bar{z}}{SE},$ $p=2,\left( 1-\Phi\left( \left| Z \right| \right) \right) ,$ $r_{\text{pooled}}=\tanh\left( \bar{z} \right)$

For longitudinal analyses, Δ-clinical scores (differences between visits) were correlated with the corresponding Z-diff values in the matched longitudinal samples. Within the clinical domain (social, sensory, adaptive, cognitive, and autism-core), tract-wise p-values were corrected using Benjamini–Hochberg FDR, reporting both nominal (p < 0.05) and FDR-adjusted (q < 0.05) results.

##### Spearman Robustness Check

Because several clinical measures showed skewed or partly bimodal distributions (especially ADOS scores), we repeated the Wave 1 correlations using Spearman’s rank (23) correlation, which is less sensitive to non-normality and outliers than Pearsons correlation. FDR correction was applied within each clinical variable, and the pooled Spearman results were compared with the Pearson estimates.

Spearman identified more FDR-significant associations than Pearson (57 vs 33), which is expected given its robustness to non-linear and non-normal relationships (24). However, there was substantial overlap between the two approaches (28 associations significant under both; 84.8% overlap relative to the larger set), indicating that the main pattern of results was not driven by distributional assumptions and supporting the use of Pearson correlations in the primary analyses.

| CROSS-SECTIONAL | | | | |
| --- | --- | --- | --- | --- |
|  | **Welch t sig tract(s)** | **Welch t score** | **Welch t nom p** | **Welch t FDR q** |
| wave 1 | — | — | — | — |
| wave 2 | — | — | — | — |
| wave 3 | Cerebral peduncle R,  Cerebral peduncle L,  External capsule R | 2.297; 2.457; −2.003 | 0.02571; 0.017; 0.04855 | 0.5635; 0.5635; 0.5635 |
|  | **MWU sig tract(s)** | **MWU score** | **MWU nom p** | **MWU FDR q** |
| wave 1 | Cingulum-hippocampus R | 10314 | 0.04433 | 0.9216 |
| wave 2 | — | — | — | — |
| wave 3 | Genu of corpus callosum,  Splenium of corpus callosum, Cerebral peduncle R,  Cerebral peduncle L,  Anterior limb of internal capsule L | 3531; 3595; 3632; 3634; 3500 | 0.03548; 0.02054; 0.0147; 0.01443; 0.04556 | 0.4258; 0.3287; 0.3287; 0.3287; 0.4374 |
| LONGITUDINAL | | | | |
|  | **Welch t sig tract(s)** | **Welch t score** | **Welch t nom p** | **Welch t FDR q** |
| wave 1–2 | — | — | — | — |
| wave 1–3 | Splenium of corpus callosum | 2.255 | 0.02891 | 0.8099 |
| wave 2–3 | Cerebral peduncle R,  Cerebral peduncle L,  Posterior corona radiata L | 2.071; 2.582; 2.047 | 0.04451; 0.0131; 0.04582 | 0.7135; 0.6288; 0.7135 |
|  | **MWU sig tract(s)** | **MWU score** | **MWU nom p** | **MWU FDR q** |
| wave 1–2 | Tapetum L | 1582 | 0.04785 | 0.9925 |
| wave 1–3 | Splenium of corpus callosum, Retrolenticular part of internal capsule R | 1266 1286 | 0.01711; 0.01097 | 0.4106; 0.4106 |
| wave 2–3 | Cerebral peduncle L | 1037 | 0.009165 | 0.4399 |

### Supplementary Table 2: Group Comparison Results

| CCA Cross-sectional results | | | | |
| --- | --- | --- | --- | --- |
| Wave | **Component** | **Mean r** | **SD r** | **Permutation p** |
| 1 | C1 | 0.264 | 0.105 | 0.001 |
|  | C2 | 0.232 | 0.163 | 0.002 |
|  | C3 | 0.188 | 0.094 | 0.001 |
| 2 | C1 | 0.324 | 0.100 | 0.001 |
|  | C2 | 0.252 | 0.133 | 0.001 |
|  | C3 | 0.183 | 0.094 | 0.002 |
| 3 | C1 | 0.031 | 0.136 | 0.330 |
|  | C2 | 0.055 | 0.129 | 0.207 |
|  | C3 | 0.037 | 0.113 | 0.299 |
| CCA Longitudinal results | | | | |
| Pair | **Component** | **Mean r** | **SD r** | **Permutation p** |
| V1-V2 | C1 | -0.122 | 0.146 | 0.867 |
|  | C2 | -0.067 | 0.097 | 0.764 |
|  | C3 | 0.011 | 0.102 | 0.434 |
| V2-V3 | C1 | 0.160 | 0.236 | 0.100 |
|  | C2 | 0.136 | 0.206 | 0.087 |
|  | C3 | 0.110 | 0.230 | 0.128 |
| V1-V3 | C1 | 0.361 | 0.173 | 0.001 |
|  | C2 | 0.224 | 0.277 | 0.014 |
|  | C3 | 0.167 | 0.271 | 0.034 |

### Supplementary Table 3: CCA Results

### Supplement Abbreviations

**ADOS** – Autism Diagnostic Observation Schedule
**ASD** – Autism Spectrum Disorder
**BLR** – Bayesian Linear Regression
**C1, C2, C3** – Canonical components 1–3 (from msCCA)
**CART** – Classification and Regression Trees
**CSS** – Calibrated Severity Score (ADOS)
**DTI** – Diffusion Tensor Imaging
**DTIFIT** – Diffusion Tensor Fitting (FSL function)
**DWI** – Diffusion-Weighted Imaging
**EPI** – Echo Planar Imaging
**EV** – Explained Variance
**FA** – Fractional Anisotropy
**FDR** – False Discovery Rate
**FOV** – Field of View
**FSIQ** – Full-Scale Intelligence Quotient
**FSL** – FMRIB Software Library
**ID** – Intellectual Disability
**ICP** – Inferior Cerebellar Peduncle
**IQR** – Interquartile Range
**JHU** – Johns Hopkins University (White Matter Atlas)
**LEAP** – Longitudinal European Autism Project
**MCP** – Middle Cerebellar Peduncle
**MP-PCA** – Marchenko–Pastur Principal Component Analysis
**MRI** – Magnetic Resonance Imaging
**msCCA** – Multi-view Sparse Canonical Correlation Analysis

**MWU**– Mann Whitney U test
**PCT** – Pontine Crossing Tract
**QC** – Quality Control
**RBS** – Repetitive Behaviour Scale
**RRB** – Restricted and Repetitive Behaviour (ADOS domain)
**SCP** – Superior Cerebellar Peduncle
**SD** – Standard Deviation
**SH** – Spherical Harmonics
**SFOF** – Superior Fronto-Occipital Fasciculus
**SLF** – Superior Longitudinal Fasciculus
**SNR** – Signal-to-Noise Ratio
**SRS** – Social Responsiveness Scale
**SSP** – Short Sensory Profile
**TBSS** – Tract-Based Spatial Statistics
**TD** – Typically Developing
**TE** – Echo Time
**TR** – Repetition Time
**UF** – Uncinate Fasciculus
**VBM** – Voxel-Based Morphometry
**VABS** – Vineland Adaptive Behaviour Scales
**WM** – White Matter
